# Structure-Function Relationships of the Vertebral Endplate

**DOI:** 10.1101/2021.05.14.444250

**Authors:** Yuanqiao Wu, Johnfredy Loaiza, Rohin Banerji, Olivia Blouin, Elise F. Morgan

**Affiliations:** Department of Mechanical Engineering, Boston University, 110 Cummington Mall, Boston, MA 02215; Department of Biomedical Engineering, Boston University, 44 Cummington Mall, Boston, MA 02215

**Keywords:** Vertebral Endplate, Micro-Computed Tomography, Bending, Density, Fracture

## Abstract

**Background:** Although deformation and fracture of the vertebral endplate have been implicated in spinal conditions such as vertebral fracture and disc degeneration, few biomechanical studies of this structure are available. The goal of this study was to quantify the mechanical behavior of the vertebral endplate.

**Methods:** Eight-five rectangular specimens were dissected from the superior and/or inferior central endplates of human lumbar spine segments L1-L4. Micro-computed tomography (μCT) imaging, four-point-bend testing, and ashing were performed to quantify the apparent elastic modulus and yield stress (modulus and yield stress, respectively, of the porous vertebral endplate), tissue yield stress (yield stress of the tissue of the vertebral endplate, excluding pores), ultimate strain, fracture strain, bone volume fraction (BV/TV), bone mineral density (BMD), and various measures of tissue density and composition (tissue mineral density, ash fraction, and ash density). Regression was used to assess the dependence of mechanical properties on density and composition.

**Results:** Wide variations in elastic and failure properties, and in density and tissue composition, were observed. BMD and BV/TV were good predictors of many of the apparent-level mechanical properties, including modulus, yield stress, and in the case of the inferior vertebral endplate, failure strains. Similar values of the mechanical properties were noted between superior and inferior vertebral endplates. In contrast to the dependence of apparent stiffness and strength on BMD and BV/TV, none of the mechanical properties depended on any of the tissue-level density measurements.

**Conclusion:** The dependence of many of the mechanical properties of the vertebral endplate on BV/TV and BMD suggests possibilities for non-invasive assessment of how this region of the spine behaves during habitual and injurious loading. Further study of the non-mineral components of the endplate tissue is required to understand how the composition of this tissue may influence the overall mechanical behavior of the vertebral endplate.

## 1. INTRODUCTION

The vertebral endplate, a thin, porous structure at the interface between the intervertebral disc and the trabecular centrum of the vertebral body, has been implicated in the etiology of two common causes of back pain, disc degeneration and vertebral fracture^1,2^. The vertebral endplate mediates fluid transport and load transfer between the disc and centrum of vertebral body, thereby serving an important structural and biochemical role in the spine^3^. Vertebral fractures, which affect at least 12-20% of men and women over the age of 50^4–6^, frequently occur at or near the vertebral endplate^7–9^. Moreover, breakage of the vertebral endplate during vertebral fracture may lead to worsening of the fracture over time^10^ and degeneration of the adjacent intervertebral disc^11–13^. Thus, study of the mechanical behavior of the vertebral endplate and the dependence of this behavior on structure and composition can aid in understanding the development and consequences of vertebral fracture.

A limited amount of data is available on the mechanical properties of the tissue in the vertebral endplate (“tissue-level properties”^14^), and less is known about the mechanical behavior of the vertebral endplate as a structure (“apparent-level properties”). Previous studies have carried out micro-indentation tests on tissue from the vertebral endplate, vertebral trabecular bone, and the cortical shell, and have found similar elastic moduli among these three types of tissue^15,16^. However, given that the vertebral endplate has a preponderance of microscale pores, mechanical characterization at larger length scales is still needed. Several studies have used much larger indenters (3 mm and 1.5 mm) to indent across the superior and/or inferior endplate surface of the vertebra. These studies have generally found that the ring apophysis is stronger and stiffer than the central region^17–21^, although the opposite was found when the cartilage endplate was left attached to the vertebral endplate^22^. It is important to note that these macro-level indentation tests do not measure the properties of the vertebral endplate alone but rather those of the vertebral endplate together with some fraction of the rest of the vertebra. The indentation strength and stiffness measured in these types of macroscale tests are lower upon removal of the vertebral endplate^23^, which adds to the evidence of the mechanical importance of the vertebral endplate but does not provide direct quantification of its properties.

Despite the paucity of mechanical data on the vertebral endplate, data on its microstructure and composition suggest that its mechanical behavior may vary greatly. Porosity and thickness tend to be higher and lower, respectively, in the superior (relative to the vertebra) vertebral endplate compared to the inferior one^24,25^, which is consistent with clinical observations of a higher incidence of fractures in the superior half of the vertebral body^26^. Among individuals, variations in the bone mineral density (BMD) of the vertebral endplate are nearly as large as those in the BMD of the entire vertebral body and in the BMD of vertebral trabecular bone^27^. Some of this variation could be due to changes in porosity; for example, both Rodriguez *et al*. and Zehra *et al*. found that vertebral endplate porosity increases approximately two-fold over the course of disc degeneration^28,29^. However, conflicting reports exist as to whether porosity and BMD change with age^30^, and the implications of the inter-and intra-individual variations in microstructure and composition for mechanical behavior are not yet known.

As such, the overall goal of this project was to characterize the mechanical behavior of the vertebral endplate. Rectangular specimens of the vertebral endplate were subjected to four-point bend tests and underwent microstructural and compositional analyses. Our specific objectives were: (1) to quantify the elastic, yield and fracture properties of superior and inferior vertebral endplates; and (2) to determine the dependence of these properties on measures of structure and composition.

## 2. MATERIALS AND METHODS

### 2.1 Specimen preparation

L1-L4 vertebrae were obtained from 39 fresh frozen cadavers (24 males, 15 females) of mean age 77.7 years (stdev = 6.5 years, range: 25-91 years) (Figure 1). The vertebral bodies were separated into superior and inferior halves with an autopsy saw, and on each half the cartilage endplate was removed with a scalpel to expose the vertebral endplate. The halves of the vertebral bodies were further trimmed using a diamond wafering blade (IsoMet 4000; Buehler, Lake Bluff, Illinois, USA) to produce a rectangular test specimen of the vertebral endplate with approximate dimensions 30mm×13mm×1.5mm (Figure 2A). Due to the irregular thickness and surface topography of the vertebral endplate, the test specimens contained some struts of subchondral trabecular bone (Figure 2B). Five of the vertebral endplates produced two test specimens each, while the remaining 80 produced only one test specimen each.

**FIGURE 1.**
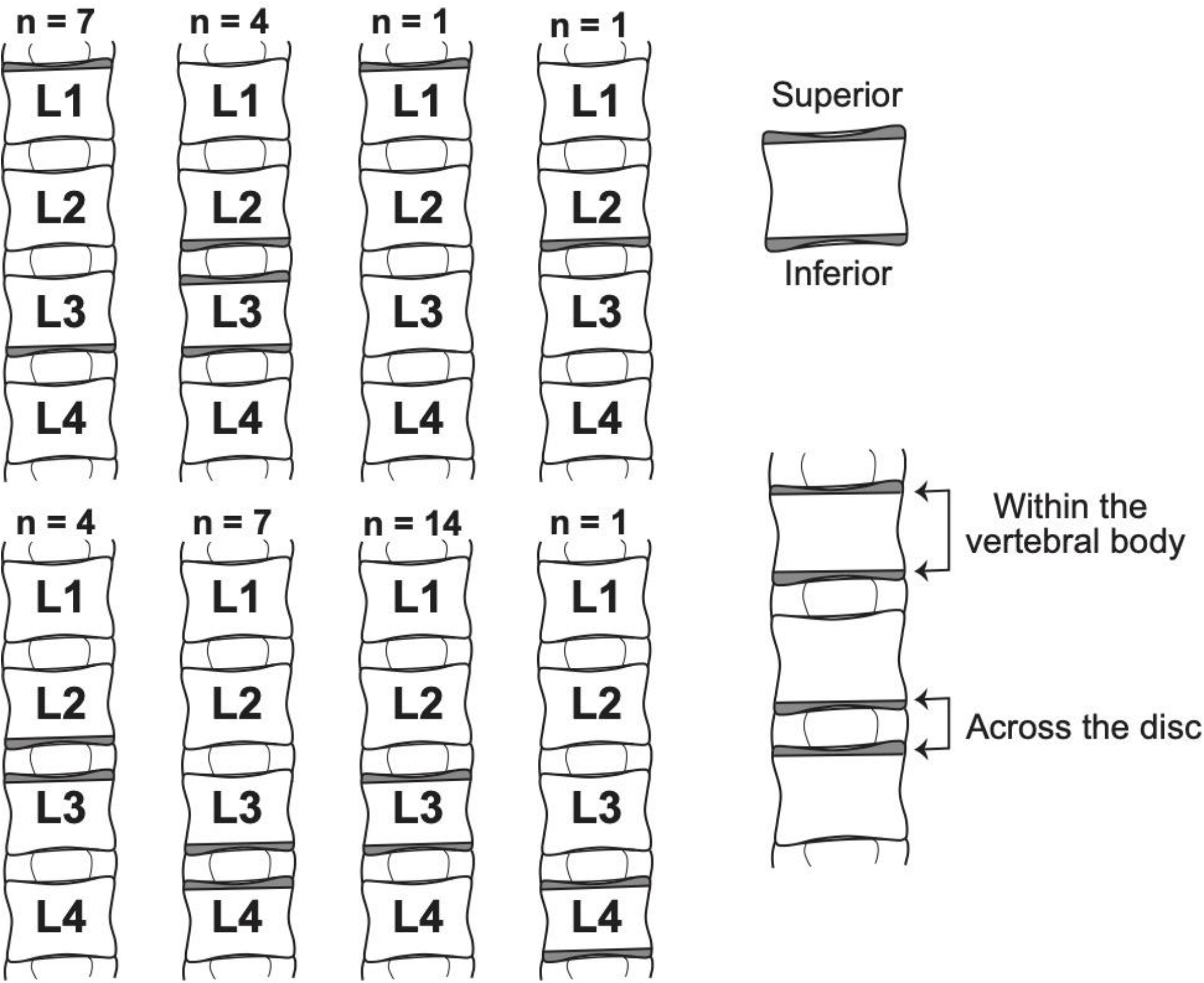
Origin of the vertebral endplate specimens used in this study with respect to the 39 L1-L4 spine segments. Gray shading indicates where the specimens were harvested from. n is the number of spine segments in each dissection scenario. Superior and inferior endplates collected from the same spine can be paired within the same vertebral body or across the same disc

**FIGURE 2.**
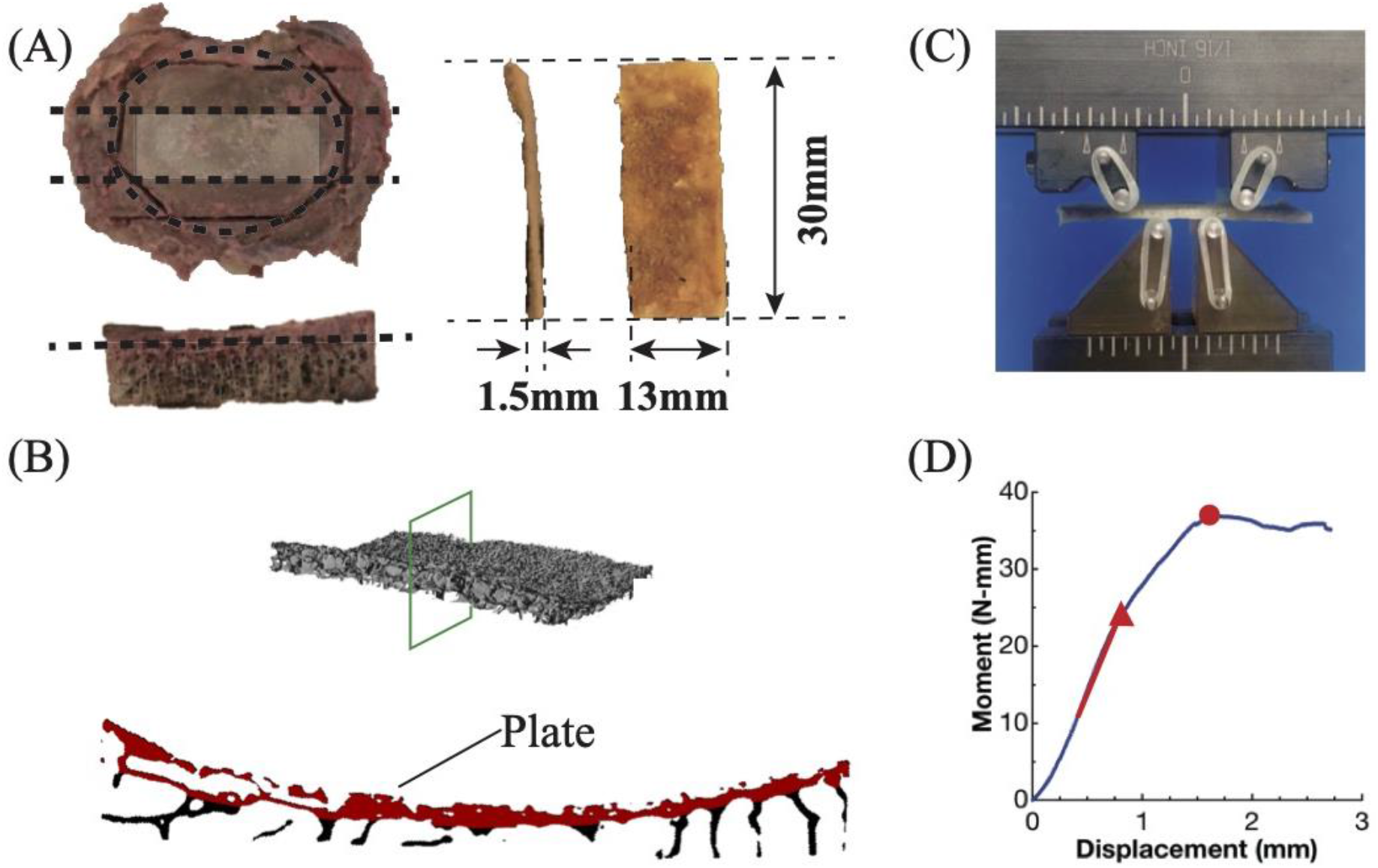
A, Specimen preparation (dotted lines represent cuts): The ring apophysis of the superior or inferior half of the vertebra was first trimmed off using bone saw followed by a transverse cut to reduce the thickness to 5mm. Then two sagittal cuts were made to dissect the central 13 mm region. The last cut further reduced the thickness to 1.5mm. B, A μCT cross-section of the vertebral endplate. The plate itself is false-colored red. C, Four-point-bend test set-up. The bottom two pins are loading pins with inner span of 8mm. The top two pins are supporting pins with outer span of 16mm. D, Representative moment-displacement curve. The red line, red triangle and red dot mark the slope used to compute the elastic modulus, yield point and ultimate point respectively

### 2.2 Micro-Computed Tomography (μCT) scanning

Each test specimen was submerged in PBS solution and imaged in a μCT scanner (μCT 40; Scanco Medical, Brüttisellen, Switzerland, 16μm/voxel, 70kV, 114μA). A threshold of 510 mg HA/cm^3^ (215 per-mille), determined from an adaptive, iterative technique (Scanco Medical), was used to binarize those μCT images. Bone volume fraction (BV/TV), bone mineral density (BMD), and tissue mineral density (TMD) were quantified for the central 16 mm of the specimens; this region corresponds to the flexural span in the bend tests. BMD was defined as the average density of all voxels in the 16mm span and is akin to apparent density, whereas TMD was defined as the average density of only the voxels within the 16mm span whose mineral density was above the threshold. Plate thickness (Figure 2B) was measured using the 3-D thickness measurement algorithm in BoneJ^31^, which computes local values of thickness throughout the structure and then averages these. Although some of the local thicknesses corresponded to the trabecular struts, their effect on the resulting average was small due to the small percentage of struts present.

### 2.3 Mechanical testing

Following μCT scanning, each specimen was placed on the support pins of a four-point-bend test fixture (inner span = 8 mm, outer span = 16 mm) in an electromechanical test frame (model 5565; Instron, Norwood, Massachusetts, USA) (Figure 2C). Specimens were oriented such that the surface of the vertebral endplate was placed in compression during the bend test, to mimic the type of concavity that develops in most clinical vertebral fractures in the elderly. After 15 cycles of preconditioning to 0.75 mm, each specimen was loaded to failure at a rate of 0.21 mm/sec. Force and displacement were measured with a 1kN load cell and the test frame’s LVDT, respectively; the measured displacement was that of the outer pins. The test was stopped when either failure of specimen occurred (defined as the force dropping to zero) or when the displacement limit of the test was reached (defined as the onset of pinching of the specimen between the upper pins and the sides of the bottom fixture, Figure 3A).

**FIGURE 3.**
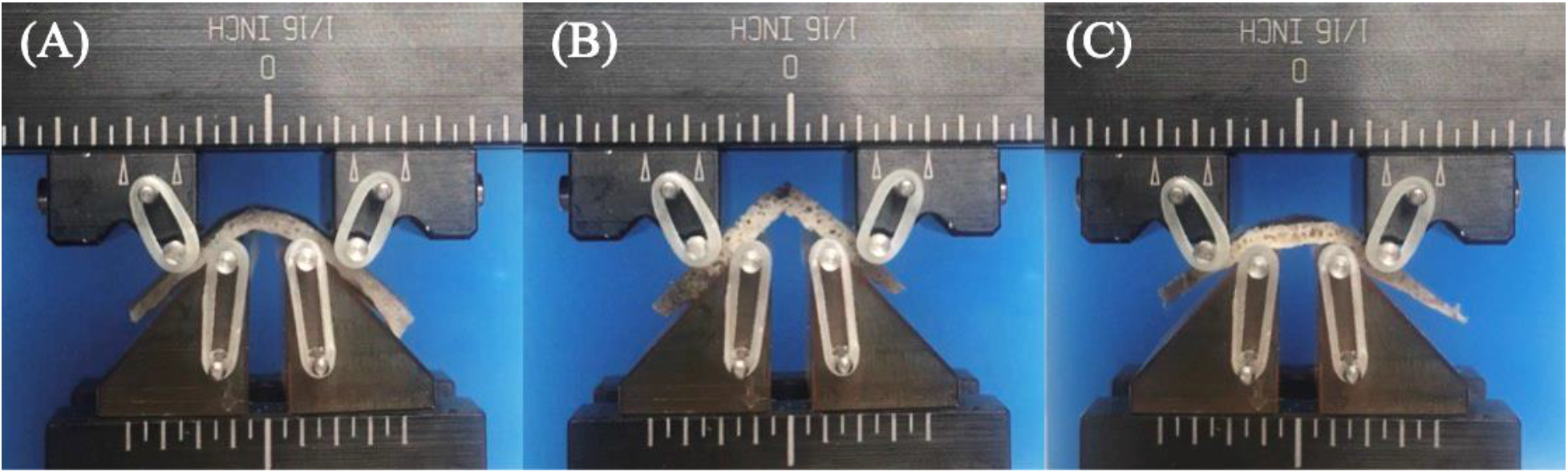
Vertebral endplates exhibited different modes of deformation and failure: A, Uniform curvature across the bending span; B, Breakage; and C, Non-uniform curvature across the bending span. All images correspond to the end of the test. The image in A illustrates the displacement limit of the test, as further applied displacement would result in pinching of the specimen between the upper pins and bottom fixture.

The apparent modulus and apparent yield stress were computed using linear elastic beam theory:

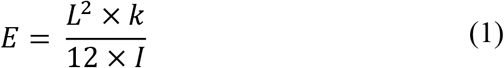

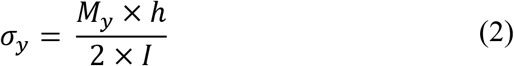

where k is the maximum slope of the moment-displacement curve prior to any local maximum in the curve (red line in Figure 2D), *M_y_* is the moment just after this region of maximum slope (red triangle in Figure), L=16 mm, *I* = 1/(12 × *b* × *h^3^*), and *b* and *h* are the width and thickness of the central 16 mm of the specimen. The tissue-level yield stress (*σ_tissue_*) was calculated at the apparent yield point, using *M_y_/S_min_*, where *S_min_* is the minimum section modulus and was computed according to published methods^32,33^ by considering all bone voxels (voxels above the threshold) within the 16mm span in the μCT image.

Video taken of the mechanical tests was synchronized to the loading curve and used to compute ultimate and fracture strains. In the still frame corresponding to the ultimate point, the curvatures of upper and lower boundaries of the 16mm span of the specimen were determined using edge detection and best-fit circles (MATLAB) and then averaged. Ultimate strain (*ε*_u_) was calculated as the distance between two edges divided by the average of the two circles’ radii. For 25 specimens, the fracture strain (*ε*_f_) was calculated using the same method for calculating the *ε*_u_, except at the point in the test when an audible cracking noise occurred at the same time as a sudden drop bending moment. For some specimens, the fracture point coincided with the ultimate point, whereas for others, it occurred after the ultimate point. Due to the displacement limit of the test, the ultimate strain and fracture strain were not obtainable for all the specimens.

### 2.4 Measurements of tissue and ash densities

After mechanical testing, each specimen was gently cleaned with a water jet to remove the bone marrow and then weighed (Mettler AT 200; Mettler Toledo, Columbus, Ohio, USA) in air and again in degassed water in order to calculate the tissue density (*ρ*_tissue_) using Archimedes’ principle. Specimens were then defatted in acetone for 12 hours and cut in half, and one half was retained for measurement of ash density. The tissue volume of this half was also calculated using Archimedes’ principle and was then dried in a muffle furnace (Thermolyne Furnace 47900, Thermo Fisher Scientific, Waltham, MA, USA) at 110°C for two hours to obtain the dry weight. The dried specimen was then put back to the furnace for another 14 hours under 650°C to obtain the ash weight. Ash fraction (*p*) and ash density (*ρ*_ash_) were computed as the ratio of ash weight to dry weight and ash weight to tissue volume, respectively. Finally, the ash weight, water weight and organic weight were each computed as a percentage of the tissue weight.

### 2.5 Statistical analyses

In cases where two specimens were obtained from the same half vertebral body, the mean value over the two specimens was used in the statistical analyses. All properties except TMD, *p, ε*_u_ and *ε*_f_ were log-transformed to follow the normal distribution assumption of statistical models. Linear regression (JMP, SAS Institute) analysis was used to determine the dependence of: 1) mechanical properties on density and mineral content, and 2) properties on age. In accordance with Hernandez et al.^34^, multiple regression analysis was also performed to test the dependence of apparent modulus and strength on both BV/TV and ash fraction. Pearson correlation analysis was used to describe the association between mechanical properties. Since properties of the vertebral endplates that come from the same donor are not independent from each other, all of the above statistical analyses were performed separately for superior and inferior vertebral endplates separately. Subsequently, paired t-tests were used to identify differences in properties between superior and inferior. These comparisons were performed for endplates spanning the same intervertebral disc (i.e. L3 superior vs. L2 inferior)^29^ and, separately, for endplates spanning the same vertebral centrum (i.e. L3 superior vs. L3 inferior)^24,25^ (Figure 1). For more general comparison of the difference between locations, nonparametric Wilcoxon test was used (and with no log transformation). A significance level of 5% was used in all analyses, and any results that resulted from overly influential data points were excluded.

## 3. RESULTS

Wide variations in vertebral endplate mechanical behavior were observed among specimens. For most of the mechanical properties, values ranged more than ten-fold (Table 1). These large ranges were observed for specimens from both superior and inferior endplates, and few differences in properties between these two locations were found (Table 1). Different modes of deformation were also observed (Figure 3). Some specimens remained intact, with no visible fractures at the end of the test (Figure 3A, C), while others broke into two parts before reaching the displacement limit (Figure 3B). Different deformation scenarios were detected among the specimens in the former category: whereas some exhibited uniform curvatures across the 16mm loading span (Figure 3A) others did not (Figure 3C).

**TABLE 1.** Properties of superior and inferior vertebral endplate specimens tested in this study. Data are presented as mean ± S.D (minimum, maximum)

|  | Location |  |
| --- | --- | --- |
|  | Superior | Inferior |
| **Plate Thickness (mm)** | 0.225 ± 0.073 (0.138, 0.457) | 0.265 ± 0.092 (0.134, 0.509) |
| **$\rho_{\text{tissue}}$ (g/cm<sup>3</sup>)** | 1.519 ± 0.107 (1.291, 1.753) | 1.498 ± 0.107 (1.265, 1.696) |
| **BV/TV (-)<sup>a,c</sup>** | 0.246 ± 0.07(0.133, 0.432) | 0.296 ± 0.109 (0.118, 0.629) |
| **BMD (g HA/cm<sup>3</sup>)<sup>a,c</sup>** | 0.286 ± 0.074 (0.168, 0.484) | 0.337 ± 0.114 (0.132, 0.641) |
| **TMD (g HA/cm<sup>3</sup>)** | 0.998 ± 0.029 (0.925, 1.072) | 1.002 ± 0.028 (0.937, 1.058) |
| **$p$ (-)** | 0.595 ± 0.084 (0.364, 0.779) | 0.607 ± 0.097 (0.369, 0.847) |
| **$\rho_{\text{ash}}$ (g/cm<sup>3</sup>)** | 0.780 ± 0.174 (0.413, 1.156) | 0.823 ± 0.171 (0.525, 1.354) |
| **E (MPa)** | 178 ± 158 (46.0, 713) | 208 ± 177 (15.5, 879) |
| **$\sigma_y$ (MPa)** | 3.48 ± 2.51 (0.94, 11.8) | 4.40 ± 3.45 (0.60, 14.3) |
| **$\epsilon_u$ (-)** | 0.065 ± 0.028 (0.016, 0.134) | 0.061 ± 0.027 (0.013, 0.116) |
| **$\epsilon_f$ (-)** | 0.079 ± 0.033 (0.020, 0.133) | 0.080 ± 0.019 (0.055, 0.121) |
| **$\sigma_{\text{tissue}}$ (MPa)<sup>b</sup>** | 27.49 ± 11.60 (10.05, 56.42) | 38.26 ± 19.28 (6.359, 75.47) |
| **Organic%** | 36.1% ± 8.5% (19.2%, 57.3%) | 35.3% ± 9.7% (14.3%, 61.5%) |
| **Mineral%** | 52.7% ± 8.1% (31.8%, 67.9%) | 54.2% ± 8.8% (36.0%, 79.2%) |
| **Water%** | 11.2% ± 7.6% (0.9%, 36.2%) | 10.6% ± 6.2% (2.2%, 29.7%) |

<sup>a</sup> $p < 0.05$ for superior vs. inferior vertebral endplates, according to an unpaired comparison (Wilcoxon test)
<sup>b</sup> $p < 0.05$ for superior vs. inferior endplates when compared within the same vertebral body
<sup>c</sup> $p < 0.05$ for superior vs. inferior endplates when compared across the disc

Much of the variation in apparent mechanical properties could be explained by variations in BMD and BV/TV (Table 2). Both the apparent modulus and apparent yield stress increased with increasing BMD (Figure 4A, B). The ultimate strain and fracture strain decreased with increasing BMD for inferior specimens while no dependence was seen for superior specimens (Figure 4C). Similar results were found when these four mechanical properties were regressed against BV/TV rather than BMD (Table 2). Consistent with these relationships between mechanical properties and both BMD and BV/TV, apparent modulus was positively correlated with apparent yield stress and, for inferior specimens, negatively correlated with fracture strain (r=-0.67, p=0.02) (Figure 5). None of the apparent mechanical properties were correlated with ultimate strain (Table 3) or depended on age (Table 2).

**TABLE 2.**
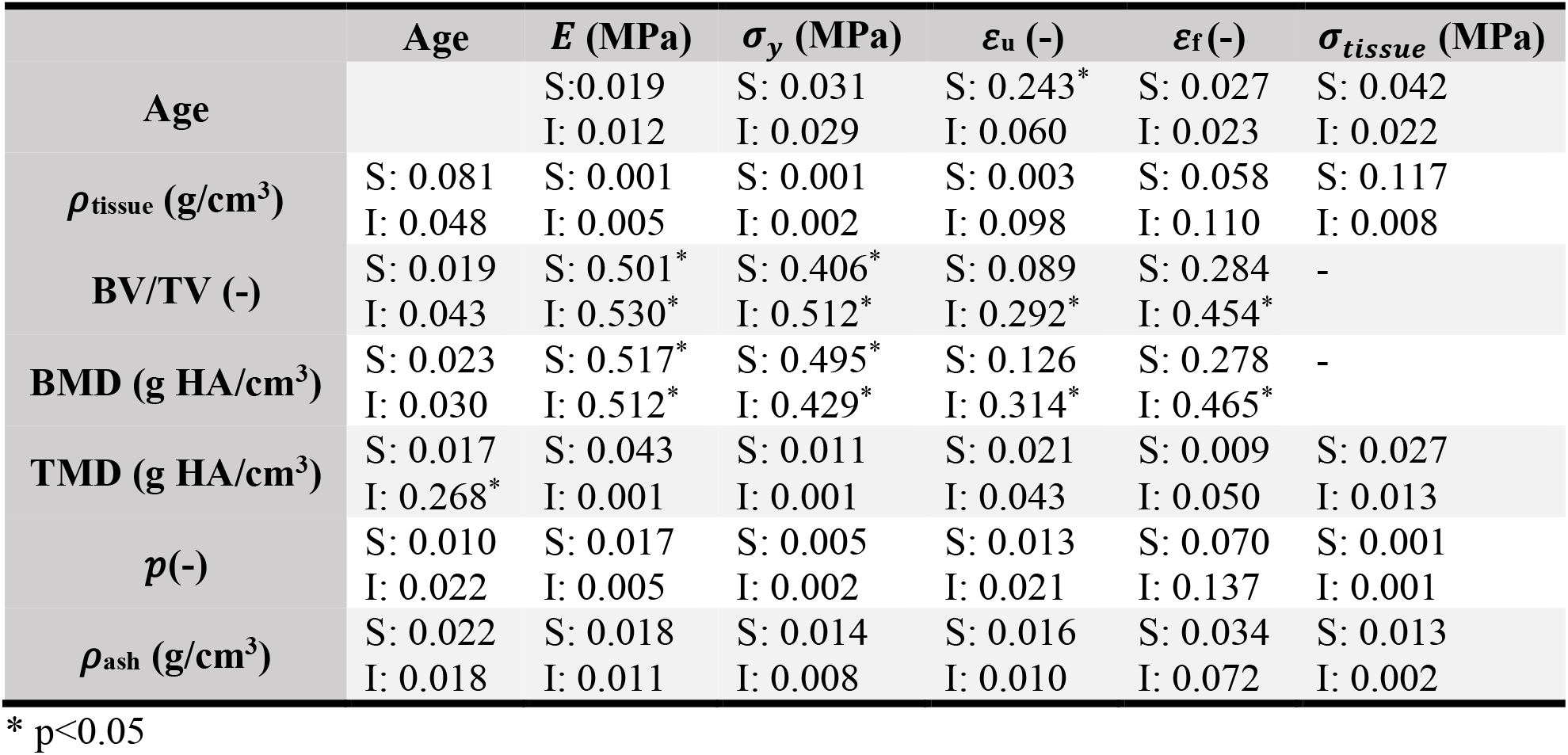
Coefficient of determination(*R*^2^) for univariate linear regressions of mechanical properties against age and measures of structure and composition. Regressions were performed through a general linear model using log transformations of the data. S and I represent the superior and inferior vertebral endplate, respectively

**FIGURE 4.**
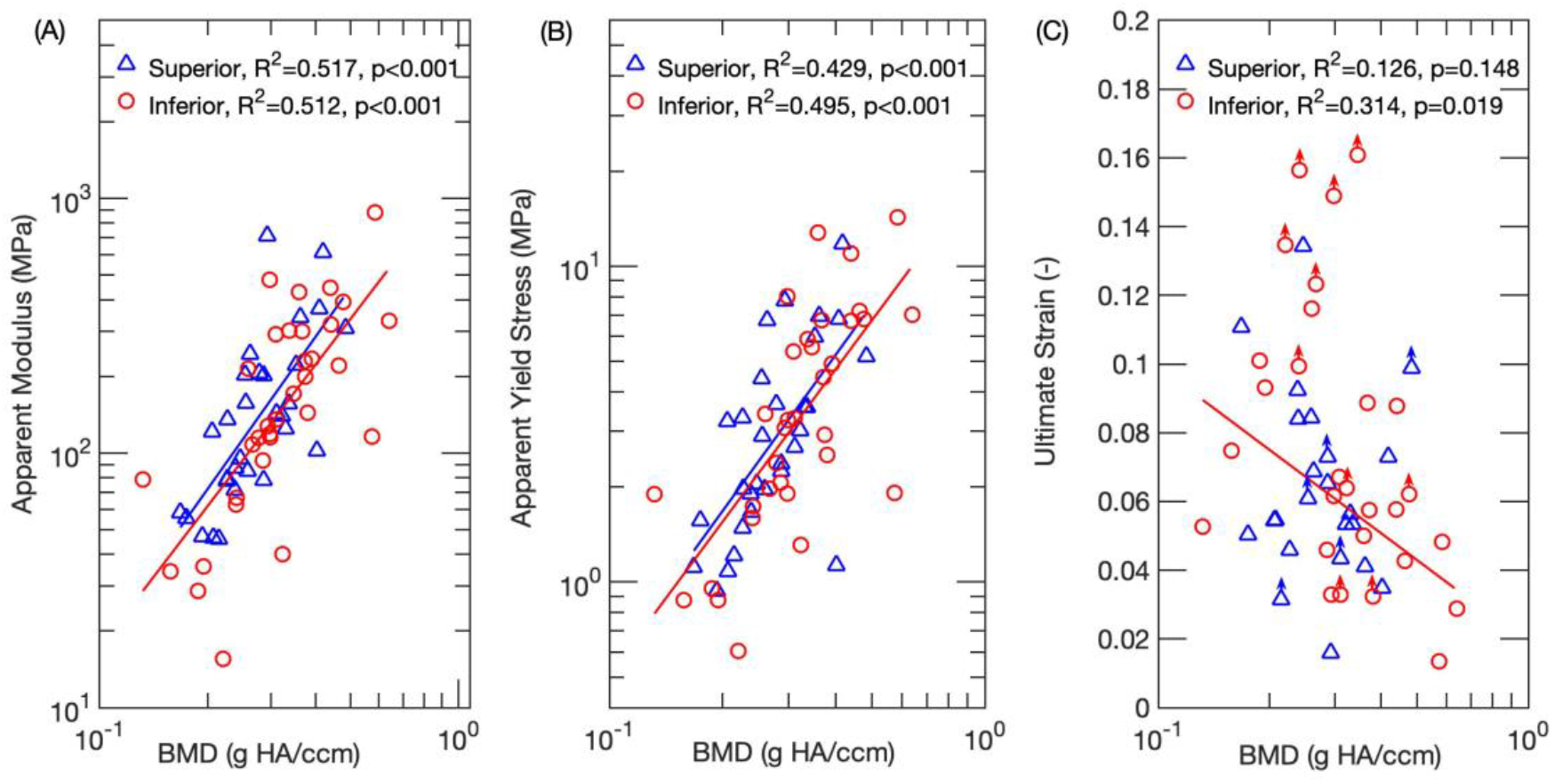
Log-log plots for A, apparent modulus; and B, apparent yield stress as a function of BMD. Regression lines are shown where applicable. Both the apparent modulus and yield stress increased with increasing BMD, for both superior and inferior endplates. C, Semi-log plot for ultimate strain as a function of BMD. An inverse relationship was found for the inferior vertebral endplates only. Points labeled with an arrow correspond to specimens that did not reach their ultimate point before the end of the test. The points are position at the largest strain that was measured in the test. These points were not included in the statistical analyses

**FIGURE 5.**
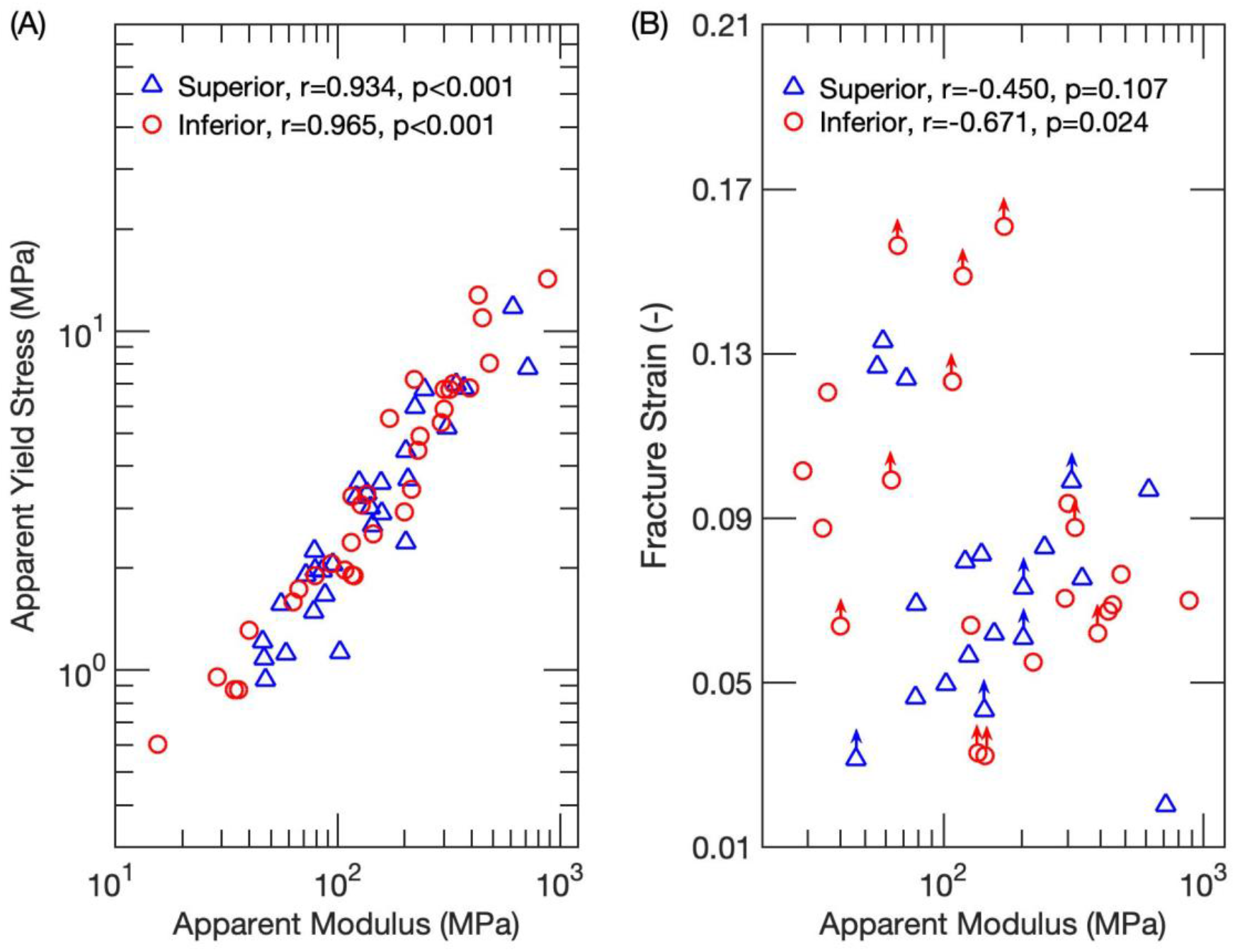
A, Log-log plots for apparent yield stress against apparent modulus for superior and inferior vertebral endplates. A positive correlation was found for both superior (r = 0.934) and inferior (r = 0.965) specimens. B, Semi-log plot for fracture strain against apparent modulus. Only fracture strain of inferior vertebral endplate was negatively correlated with apparent modulus. Points labeled with an arrow correspond to specimens that did not reach their fracture point before the end of the test. The points are position at the largest strain that was measured in the test. These points were not included in the statistical analyses

**TABLE 3.**
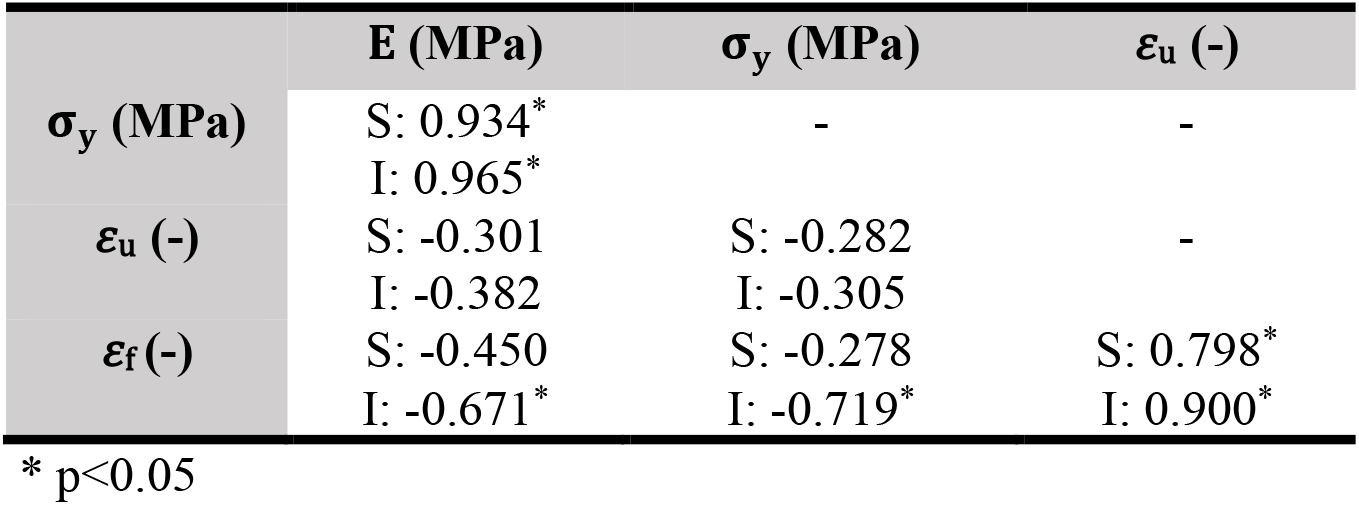
Pearson correlation (r) mechanical properties. S and I represent the superior and inferior vertebral endplate respectively

In contrast to the strong dependence of the mechanical properties of the vertebral endplate on BMD and BV/TV, little dependence on measures of tissue-level density or ash density was found. None of the mechanical properties, including tissue yield stress, depended on any of the tissue-level density measurements (Table 2). Adding ash fraction to the regression of apparent modulus and strength against BV/TV also did not improve the fit over that obtained with BV/TV alone (p > 0.091).

## 4. DISCUSSION

In light of the evidence that the vertebral endplate is a biomechanically critical structure, the goal of this study was to quantify its mechanical behavior. We found that BMD and BV/TV were strong predictors of many of the apparent-level mechanical properties, including modulus, yield stress, and in the case of the inferior vertebral endplate, failure strains. We also found that both the apparent modulus and apparent yield stress were inversely correlated with the failure strains. Similar values of the mechanical properties were noted between superior and inferior vertebral endplates, despite some small differences in thickness, BMD and BV/TV. In contrast to the strong dependence of apparent stiffness and strength on BMD and BV/TV, none of the mechanical properties depended on any of the tissue-level density measurements. These results indicated that amount of bone tissue present, rather than the composition of that tissue, is the most important determinant of the mechanical behavior of the vertebral endplate.

This study has several strengths. Principally, isolating the vertebral endplate from the rest of the vertebra enabled us to measure its properties directly. In prior studies, indentation tests were performed along the superior or inferior surfaces of the vertebral body, and thus those results correspond to the combined mechanical behavior of the endplate and underlying bone tissue^17,18,20–22^. Having isolated the vertebral endplate, we could also directly measure the tissue density, ash density and ash fraction, all of which can influence the mechanical behavior^35–38^, rather than relying on estimates of density from computed tomography scans^24,25,27–29^. The mechanical testing method used in this study has some additional advantages are that we used a physiological loading mode, flexion^10^, and a length scale on par with that of the endplate deflections and deformities associated with vertebral fracture and disc degeneration^39–42^. Finally, we tested specimens from the central endplate, rather than the ring apophysis, given the high prevalence of vertebral fractures and other types of damage that occur in this region^9,43,44^.

This study also has limitations. First, some subchondral trabecular bone was present in the specimens because of the irregular topography of the boundary between the vertebral endplate and trabecular centrum and because of the difficulty in discerning this boundary during specimen preparation. However, the μCT images revealed very few, if any connections between the short struts of trabeculae that were present, indicating that they would have minimal contribution to the flexural behavior. Second, the definition of the apparent yield stress using a decrease in slope in the moment-displacement curve involves some subjectivity. To mitigate bias, we also calculated the apparent yield stress using the “fully plastic bending moment” (Appendices). The two yield stresses were highly correlated with one another and exhibited very similar statistical results regarding the dependencies on density measures and correlations with apparent modulus, giving us confidence in the results we reported here. Third, the low flexural rigidity and appreciable ductility of many of the specimens meant that the displacement limit of the testing fixture was reached before these specimens reached their fracture strain or, in some cases, even ultimate strain. The limited number of data points reduced the power to detect differences in failure strains between superior and inferior endplates and to detect associations between the failure strains and other properties. A final limitation to report is the age of donors was not uniformly distributed. Only two donors of the 39 donors were under age 60. Although the predominance of older donors in our data set makes the results meaningful for aging-related conditions such as vertebral fracture, whether the results of this study extend to younger spines is unclear.

The lack of dependence of the measured mechanical properties on ash fraction (Table 2) was somewhat unexpected. Prior work has found associations between ash fraction and several mechanical properties, particularly stiffness and strength, of cortical bone, trabecular bone, and several other mineralized tissues^35,37,38^. Hernandez et al. found that, for femoral cortical bone and vertebral trabecular bone, predictions of both strength and stiffness using BV/TV were improved by also using ash fraction^34^. However, in our study, this was generally not the case (Appendices), even though both we and Hernandez et al. found that BV/TV explained more of the variation in strength and stiffness than did the ash fraction. This discrepancy between current and previous results may be due to the larger range of BV/TV values examined by Hernandez et al. (range: 0.02~0.84 vs. 0.118~0.629 in the current study), since they included both trabecular and cortical bone, and to differences in structure and composition between the vertebral endplate and bone in other skeletal sites. As compared to mean values reported for vertebral trabecular bone, trabecular bone in several other anatomic sites and cortical bone in the lower limb, the vertebral endplate exhibits lower tissue density and ash density, and similar ash fraction (Table 4), which altogether suggests higher water content. More detailed examination, using water, mineral, organic weight fractions, reveals a cohort of vertebral endplate specimens with high water fraction (Figure 6), but also a cohort with higher organic fraction than vertebral trabecular tissue, and overall a much larger compositional range than vertebral trabecular tissue. The vertebral endplate specimens also tended to exhibit lower mineral content than has been reported for cortical bone (Figure 6); although a comparison to the vertebral cortical shell in particular would be relevant, no such density measurements are available. These variations in aspects of composition other than ash fraction, as well as properties of the organic phase itself, such as collagen content and cross-link density^45,46^, may contribute more to the mechanical properties of the vertebral endplate than does ash fraction.

**TABLE 4.** Measures of density and ash fraction for the human vertebral endplate, human trabecular bone, and human cortical bone

| Properties | Bone Type | Anatomic Site | Mean ±STD | Range |
| --- | --- | --- | --- | --- |
| Tissue Density (g/cm<sup>3</sup>) * | Vertebral Endplate | Vertebral Body | 1.508 ± 0.107 | 1.265, 1.753 |
| | Trabecular Bone | Vertebral Body<sup>54</sup> | 1.93 ± 0.068 | NR |
| | | Vertebral Body<sup>55</sup> | 2.01 ± 0.17 | 1.76, 2.41 |
| | | Proximal Tibia<sup>55</sup> | 2.08 ± 0.05 | 1.88, 2.13 |
| | Cortical Bone | Vertebral Body<sup>56</sup> | 1.79 ± 0.04 | 1.42, 1.94 |
| | | Femoral Neck<sup>55</sup> | 2.11 ± 0.07 | 1.93, 2.20 |
| | | Femoral Diaphysis<sup>57</sup> | 1.84 ± 0.13 | 1.51, 2.00 |
| | | Femoral Diaphysis<sup>58</sup> | 1.88 ± 0.05 | NR |
| | | Greater Trochanter<sup>55</sup> | 2.09 ± 0.03 | 2.02, 2.15 |
| | | Tibial Diaphysis<sup>58</sup> | 1.80 ± 0.19 | NR |
| | | Tibial Diaphysis<sup>59</sup> | 1.86 ± 0.06 | 1.75, 1.95 |
| BV/TV (-) | Vertebral Endplate | Vertebral Body | 0.272 ± 0.096 | 0.118, 0.629 |
| | | Vertebral Body<sup>29</sup> | 0.36 ± 0.13 | 0.10, 0.68 |
| | Trabecular Bone | Vertebral Body<sup>56</sup> | 0.13 ± 0.01 | 0.05, 0.27 |
| | | Vertebral Body<sup>60</sup> | 0.18 ± 0.06 | 0.05, 0.27 |
| | | Vertebral Body<sup>55</sup> | 0.09 ± 0.02 | 0.04, 0.18 |
| | | Proximal Tibia<sup>55</sup> | 0.11 ± 0.04 | 0.05, 0.20 |
| | | Femoral Neck<sup>55</sup> | 0.27 ± 0.07 | 0.12, 0.36 |
| | | Greater Trochanter<sup>55</sup> | 0.11 ± 0.02 | 0.07, 0.14 |
| | | Distal Tibia<sup>61</sup> | 0.15 ± 0.03 | NR |
| | Cortical bone | Distal Tibia<sup>61</sup> | 0.88 ± 0.04 | NR |
| | | Femoral Neck<sup>62</sup> | 0.89 ± 0.09 | 0.6, 0.98 |
| | | Femoral Neck<sup>63</sup> | 0.92 ± 0.03 | 0.84, 0.96 |
| | | Femoral Diaphysis<sup>64</sup> | 0.83 ± 0.11 | 0.24, 0.98 |
| | | Diaphysis from both Tibia and femur<sup>65</sup> | 0.83 ± 0.13 | 0.54, 0.97 |

| Properties | Bone | Anatomic Site | Mean ±STD | Range |
| --- | --- | --- | --- | --- |
| Ash Fraction<br>(-) | Vertebral Endplate | Vertebral Body | $0.601 \pm 0.090$ | 0.364, 0.847 |
| | Trabecular Bone | Vertebral Body<sup>56</sup> | $0.64 \pm 0.02$ | 0.59, 0.69 |
| | | Vertebral Body<sup>36</sup> | $0.61 \pm 0.02$ | 0.53, 0.66 |
| | Cortical Bone | Femoral Diaphysis<sup>36</sup> | $0.58 \pm 0.10$ | 0.17, 0.66 |
| | | Femoral Diaphysis<sup>35</sup> | $0.65 \pm 0.02$ | NR |
| | | Femoral Diaphysis<sup>58</sup> | $0.64 \pm 0.01$ | NR |
| | | Tibial Diaphysis<sup>58</sup> | $0.63 \pm 0.02$ | NR |
| | | Tibial Diaphysis<sup>59</sup> | $0.66 \pm 0.02$ | 0.61, 0.69 |
| Ash Density<br>(g/cm<sup>3</sup>) | Vertebral Endplate | Vertebral Body | $0.802 \pm 0.173$ | 0.413, 1.354 |
| | Trabecular Bone | Vertebral Body<sup>56</sup> | $1.01 \pm 0.03$ | 0.78, 1.17 |
| | Cortical Bone | Femoral Diaphysis<sup>57</sup> | $1.10 \pm 0.08$ | 0.9, 1.21 |
| | | Femoral Diaphysis<sup>66</sup> | $1.05 \pm 0.14$ | 0.64, 1.20 |
| | | Diaphysis from both Tibia and femur<sup>65</sup> | $0.99 \pm 0.13$ | 0.68, 1.22 |
| | | Children Femoral shaft<sup>67</sup> | $0.92 \pm 0.15$ | 0.58, 1.17 |
| | | Adult femoral and tibial shaft<sup>67</sup> | $1.15 \pm 0.07$ | 1.00, 1.29 |

\*For cortical bone, the specimen volume used to calculate tissue density and ash density
included pore space corresponding to vascular porosity as well as lacunar-canalicular porosity
NR: not reported by the study

**FIGURE 6.**
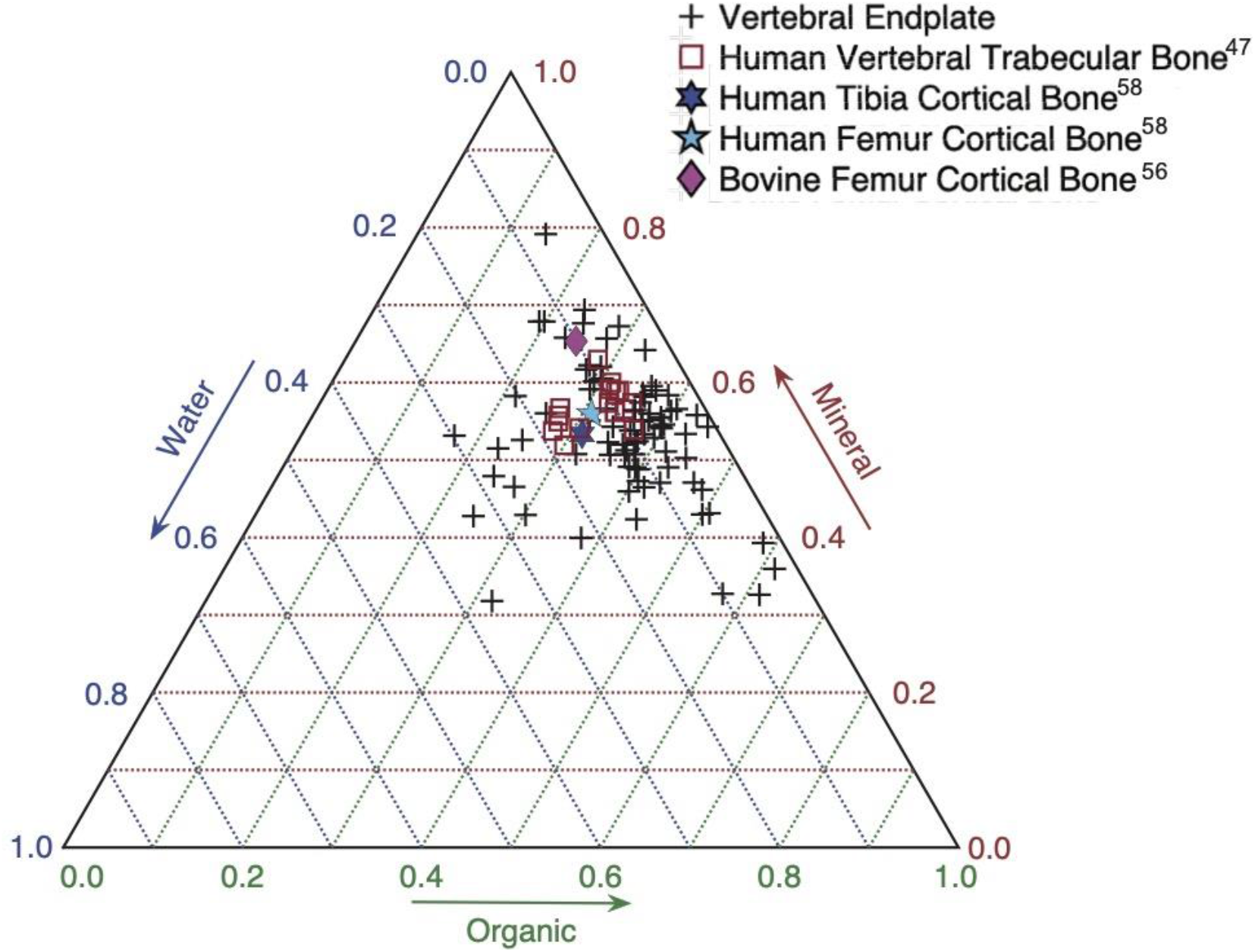
Ternary plot of the mineral, organic and water weight fractions

The inverse trend between fracture strain and each of apparent modulus and yield stress suggests a potential compensatory mechanism in the vertebral endplate. With increased BV/TV and BMD, specimens from the inferior endplates exhibited higher stiffness and strength but lower fracture strain. This tradeoff has been found for cortical bone from different anatomic sites and species^38,47^, and it indicates that specimens that are more compliant and yield at lower stresses also tend to be able to sustain greater deformation before fracturing. This tradeoff may be particularly relevant in the vertebral endplate since it is the structural boundary between the intervertebral disc and the hematopoietic tissues within the vertebral body. Fracture of this boundary has been hypothesized to trigger inflammatory cascades that can hasten degeneration of the disc^11–13^. Interestingly, the inverse trend between fracture strain and stiffness, strength, and density was not found for the superior vertebral endplates, even though the fracture strains of the superior vertebral endplate were as high as those in the inferior vertebral endplates.

Comparisons were made between superior and inferior endplates because of the suggestion that the former are more susceptible to failure^30^. Age-related vertebral fractures occur more frequently in the superior as compared to inferior half of the vertebra^26^, though whether this asymmetry is related to differences in the endplate properties themselves has not been established. Our finding of a lack of any differences in mechanical properties between superior and inferior endplates suggests that there may be other causes, such as the properties of the adjacent trabecular bone^30^, properties of the adjacent disc, and endplate curvature. Some differences in structural properties—BV/TV, BMD and thickness—were found between superior and inferior specimens but only when the comparison was made across the disc as opposed to across the vertebral body (Table 1, Figure S2). These results suggest that the structure of the vertebral endplate may be intrinsic to the vertebral body rather than the motion segment or that there may be asymmetry along the superior-inferior axis in how the vertebral endplate disc interacts with the disc.

Our findings suggest new possibilities for non-invasive assessment of vertebral fracture. Non-invasive estimation of vertebral strength and stiffness from density measurements is well established^48,49^, but thus far this approach has not been applied in a data-driven manner for the vertebral endplate. Prior work has shown that the density (*i.e*., BV/TV and BMD) of the endplate region is poorly correlated with the average density of the vertebral body, particularly in women^27^. Our data suggest that measurement of the density of the vertebral endplate can be useful for a more accurate estimate of its mechanical behavior and, by extension, the mechanisms of vertebral fracture. The wide variation of the properties of the vertebral endplate compared to that seen in trabecular and cortical bone at other anatomic sites further supports this approach. For example, the ranges of BV/TV, apparent modulus and apparent yield stress we have found for the vertebral endplate are larger than those for trabecular bone and cortical bone at single anatomic sites (Table 4)^35,50^. Using the mechanical properties obtain in the current study as input into finite element models of the vertebra may also improve the accuracy with which these models predict the where and how the vertebra deforms as it fractures^40,51^.

The wide variations in vertebral endplate properties are consistent with the growing understanding that this region of the spine is highly metabolically active. Bone remodeling, mineralization of the cartilage endplate, and inflammatory and repair responses triggered by damage to tissues in the endplate region may all result in changes in composition, density, microstructure and thickness of the vertebral endplate^29,52,53^. Bone loss due to aging and changes in the mechanical behavior of the disc due to degeneration may also affect the mechanical environment of the vertebral endplate, which in turn could result in adaptive changes in the properties of this structure. However, despite the complexity of how these myriad factors may affect a variety of physical properties of the vertebral endplate, the findings of this study indicate that only a subset of these, BV/TV and BMD, which largely describe the amount of bone tissue present, are strongly predictive of the mechanical behavior. Further study of how this subset of properties is affected by metabolic and mechanobiological demands, as well as genetic and other factors, will likely lead to improved predictions of the risks and consequences of spine injuries and pathologies.

## APPENDICES

Because of the subjectivity in determine the end of elastic regime from moment-displacement curve, a second measure of the apparent yield stress σ_u_ was calculated. The tissue of the vertebral endplate was assumed to be perfectly plastic following yield, and the maximum moment was assumed to correspond to the ‘fully-plastic bending moment’^69^. σ_u_ was calculated as:

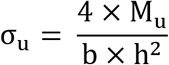

where b and h are the width and thickness of the central 16 mm of the specimen.

The mean ± STD of σ_u_ for superior and inferior vertebral endplates were 4.00 ± 2.80 (range: 1.28 ~ 11.2 MPa) and 5.35 ± 3.83 MPa (0.84 ~ 14.2 MPa) respectively. σ_u_ was highly correlated with σ_y_ (r > 0.950, p < 0.05) and, like σ_y_, σ_u_ increased with increasing BV/TV and BMD for both superior and inferior specimens (R^2^ > 0.43, p < 0.05). Adding ash fraction to the regression of σ_u_ against BV/TV was improved for inferior vertebral endplates only (R^2^ increased from 0.495 to 0.633 (p = 0.015, Figure S1).

