## Supplemental for "Structure-Function Relationships of the Vertebral Endplate"

**SUPPORTING INFORMATIONS**

To measure the plate thickness, the binarized μCT images were pre-processed with MATLAB built-in function, imfill (R2020a, MathWorks, Natick, Massachusetts, USA) to fill the pores, presented in each 2D images.

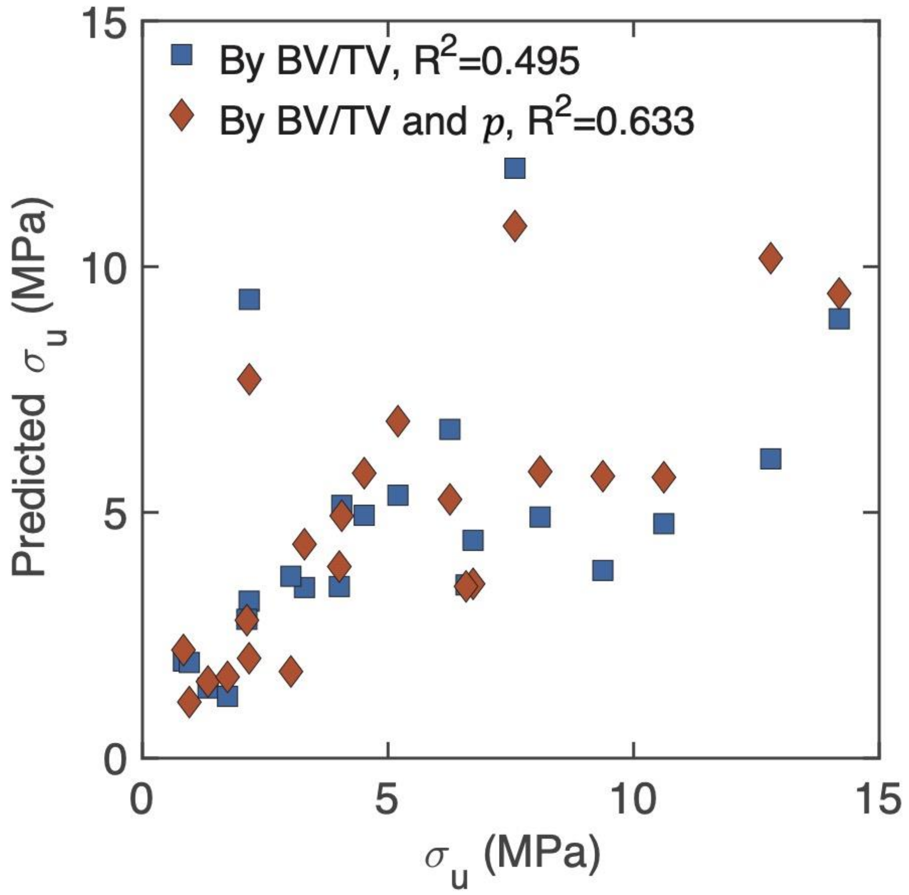


FIGURE S1, $\sigma_u$ predicted by using BV/TV only and by BV/TV together with $p$. Adding $p$ to the regression was able to explain more of the variations in $\sigma_u$

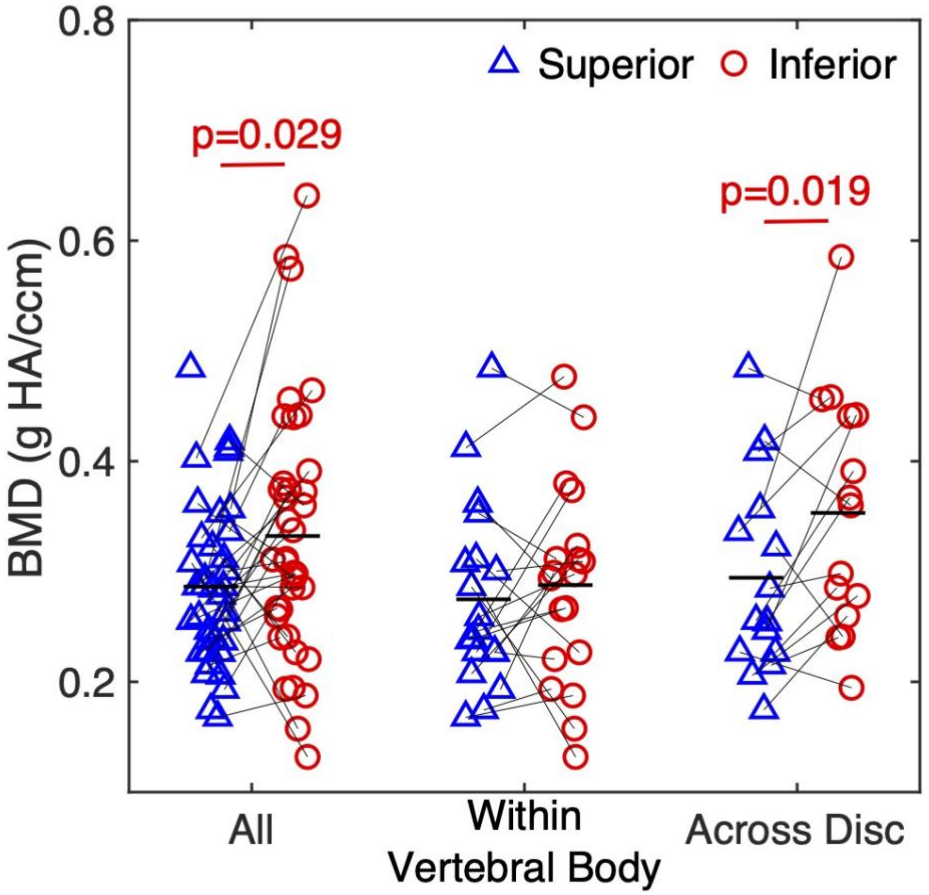


FIGURE S2, Comparison of BMD between superior and inferior vertebral endplates. When compared across the disc using a paired t-test, or when compared in an unpaired manner, BMD is higher in the inferior vertebral endplates. However, when compared within the vertebral body using a paired t-test, no difference was found
